# Improved Sleep Spindle Detection Using the BOSC Method: A Comparison with Traditional Approaches

**DOI:** 10.1101/2025.07.31.667963

**Authors:** Cameron Turnbull, Mitchell Prostebby, Ismail Babale, Silvia Pagliardini, Clayton Dickson

**Affiliations:** University of Alberta, Neuroscience and Mental Health Institute, 2-132, Li Ka Shing Centre for Health Research Innovation, 87 Avenue & 112 Street, Edmonton, AB,Canada, T6G 2E1; University of Alberta, Department of Physiology, 3020F Katz Building, 11361 87th Ave,Edmonton, AB, Canada, T6G 2E1; University of Alberta, Department of Anesthesiology and Pain Medicine, 2-150 Clinical Sciences Building 8440 112 St NW, Edmonton, Alberta T6G 2B7; University of Alberta, Department of Psychology, P217 Biological Sciences Building, 11455 Saskatchewan Drive, Edmonton, AB, Canada T6G 2E9

**Keywords:** Sleep spindles, Spindle detection, EEG, Memory consolidation, NREM sleep

## Abstract

Sleep spindles are brief bursts of 6–20 Hz local field potential (LFP/EEG) activity that occur during non-rapid eye movement sleep. Traditional spindle detection methods rely on manually set amplitude and duration thresholds, but this approach can be vulnerable to false detections and could miss low-amplitude spindles due to the inflexible nature of the thresholding technique. The Better OSCillation (BOSC) detection method offers a more robust alternative by applying frequency-specific power thresholds calibrated to the signal itself and requiring a minimum number of oscillation cycles. In this study, we compared traditional and BOSC methods for spindle detection in a variety of ways. First, we created two synthetic datasets: one with synthetic spindles modelled on previous data and one with single wave pulses that had a period consistent with the spindle frequency band. These datasets were used to demonstrate each method’s performance when given events that should be detected (synthetic spindles) and those that should not (single wave pulses). Second, we analyzed cortical local field potentials from recordings of rats during natural sleep using both methods to determine their relative effectiveness given biological LFP data. BOSC consistently outperformed the traditional approach in all situations, identifying more valid spindles while minimizing false (or likely false) detections. These findings validate BOSC as a superior method of spindle detection due to its better calibration to the actual signal.

## Intro

Sleep spindles are brief, rhythmic bursts of neocortical local field potentials (LFP)/EEG that have a bandwidth between 6–20 Hz and which occur during non-rapid eye movement (NREM) sleep. Mechanistically, spindles arise from rhythmic inhibition of thalamocortical nuclei by the reticular thalamic nucleus, which leads to rebound excitation of these excitatory neurons and their cortical targets (Steriade et al., 1993). Within the broad definition of spindles exist two subtypes: low-voltage spindles (LVS) which oscillate between 10-20 Hz (although they have been observed at as low as 6 Hz) and that have lower amplitudes, versus high-voltage spindles (HVS) which oscillate between 6-10 Hz and have higher amplitudes (Kandel and Buzsáki, 1997).

The functional relevance of HVS is unknown (Johnson et al., 2010), but LVS have been suggested to be crucial for explicit memory consolidation during sleep. Following periods of intense learning, LVS density increases during subsequent sleep epochs (Gais et al., 2002), and artificially enhancing LVS activity has been shown to improve performance on explicit memory tasks learned prior to sleeping (Marshall et al., 2006). LVS are also temporally coupled with hippocampal ripples, another local oscillatory event implicated in memory consolidation (Siapas et al., 1998). Moreover, spindle disruption in various neurological and psychiatric conditions could provide a biomarker for such disorders as schizophrenia or Alzheimer’s disease (Ferrarelli et al., 2007; Rauchs et al., 2008).

The traditional method of detecting spindles involves setting thresholds for amplitude (power) and duration, which events must exceed to be identified as spindles. To determine the power threshold, power in the spindle bandwidth is calculated across the recording and the threshold is set at a certain statistical value, commonly 1.5 times the standard deviation (SD) of the signal above the mean power level (Eckert et al., 2020). For the duration threshold, a common value for spindles is 350 ms, which is equivalent to 3 cycles of a spindle oscillation at the lower bound of its frequency range (6 Hz). Any events exceeding these thresholds are classified as spindles. However, this method may be susceptible to false detections that are not truly oscillatory and may also miss events that might otherwise be classified as spindles due to its strict dependence on thresholded power measures.

The traditional method’s reliance on the standard deviation (SD) of the signal to set power thresholds and the fact that it has not previously been probed using noisy synthetic data, limits its reliability. Power in narrow frequency bands, such as the spindle range, is often skewed and non-Gaussian, making the use of the mean and SD statistically inappropriate. Additionally, both the mean and SD of spindle-band power are easily inflated by the presence of non-oscillatory events and artifacts. This means the threshold is heavily influenced by the nature of the signal itself— including the very spindles it is intended to detect—leading to inconsistent and potentially biased detection outcomes.

The Better OSCillation detection method (BOSC), first introduced by Caplan et al. (2001) and subsequently rigorously tested by Whitten et al. (2011) and Hughes et al. (2012), derives detection thresholds in a way that is consistent and comparable across frequencies, is robust across state changes, and specifies the minimum number of cycles needed to detect any oscillation. It does so by modelling the known statistical properties of the actual signals themselves. These parameters make it an ideal method for precise spindle detection.

In the present study, the performance of these two methods was compared using both synthetic and biological LFP recordings to determine which is most effective. We report that BOSC is a superior method of spindle detection since it is robust against transient pulses and can detect significantly more spindles than the traditional method.

## Methods

### Animals

Recordings were made from 4 male Sprague Dawley rats obtained from the Health Sciences Laboratory Animal Services of the University of Alberta within a weight range of 250 - 300 g. All animals were provided with food and water ad libitum and were maintained on a 12 h light/dark cycle, with lights on at 7:00 A.M. All procedures conformed to the guidelines of the Canadian Council on Animal Care and were approved by the Health Sciences Animal Policy and Welfare Committees (AUP 461) of the University of Alberta.

### Instrumentation for Natural Sleep

In preparation for local field potential (LFP), electromyographic (EMG) and temperature probe implantation, rats were initially anesthetized with isoflurane in air (5%) followed by a maintenance concentration of 2-3% to achieve a surgical plane. Using aseptic techniques, an incision was made in the abdomen to place a temperature probe (Biomark BioTherm13 PIT Tag, Boise, ID, USA) from which readings were used for the analysis of the whole-body plethysmography data. After the implantation of the temperature probe and abdominal suturing, rats were transferred to a stereotaxic device (Kopf, Tujunga, CA, USA) and positioned to ensure that lambda and bregma were at the same horizontal level. Bipolar multi-stranded Teflon-coated stainless steel wires (AM-Systems, Inc., Carlsborg, WA, USA) with a distance between the wires of approximately 1 mm were implanted both in the neocortex (nCTX) and hippocampus (HPC) based on the following stereotaxic coordinates relative to bregma : nCTX: anterior-posterior (AP): +2.5; mediolateral (ML): −1.2; dorsoventral (DV): −1.5 to −2.0 mm; HPC: AP, −3.3; ML, −2.4; DV, −2.5 to −3.0 mm. Electrodes and the head stage pins were fixed to the skull using anchoring jeweler’s screws followed by a layer of dental acrylic. Recordings were made one week after the surgeries to allow sufficient recovery time.

### Natural Sleep and Electrophysiology Recordings

Two days prior to recording, the rats were habituated to a whole-body plethysmograph chamber (Buxco Respiratory Products, DSI, New Brighton, MN, USA) for 3 hours while also having the head pin connections for the LFP/EMG attached. A constant flow of air was delivered at a rate of 1.5 L per minute. Seven days after the initial surgeries, the rats were placed in the whole-body plethysmograph chamber from 12:00 – 14:30 while LFP and EMG signals were acquired using PowerLab (AD Instruments, Colarado Springs, CO, USA) sampled at 1 kHz. LFP signals from the nCTX and HPC were amplified at 1000 gain and filtered between 0.1 and 500 Hz using a differential amplifier (model P511, Grass Technologies., Quincy, MA, USA), whereas EMG activity was amplified at a gain of 10,000 and filtered between 100 and 500 Hz (model 1700 A-M Systems).

### Sleep State Scoring

Scoring was performed visually on LFP records using Lab Chart (AD Instruments). Wake state was identified by the presence of high neck EMG activity with low amplitude and high frequency nCTX. NREM was characterized by the presence of low muscle tone and slow-frequency high-amplitude LFP in the nCTX (delta wave: 0.5-4 Hz). REM sleep was identified by even lower muscle tone, low amplitude neocortical signals and a prominent theta rhythm from the HPC.

### Traditional Spindle Detection

Many studies that analyze spindles use the SD of mean spindle power and an arbitrary duration threshold to detect spindles (Bandarabadi et al., 2020; Eschenko et al., 2006; Siapas and Wilson, 1998), but we created an algorithm that followed the method employed by Eckert et al. (2020) except that it used continuous wavelet power instead of instantaneous power (squared voltage at each sample) for a fairer comparison with BOSC (Fig. 1 C). This algorithm first calculates the maximum wavelet power within the spindle frequency range (6-20 Hz) for each sample of the z-score normalized raw LFP (Fig. 1 A). The continuous spindle wavelet power across time was then smoothed using a 100 ms Hilbert transform to obtain the envelope of the signal. Peaks in power greater than 1.5 SD above the mean power were selected and the envelope was walked forward and backward to where the power was 0.75 SD above the mean to obtain the spindle start and end times. Only events longer than 350 ms were selected as spindles. After this preliminary spindle detection, the trough-to-trough duration of each individual cycle within each detection was measured and if the detected spindle contained any cycles of duration greater than 125 ms it was removed. Last, any detections within 0.1 seconds of each other were merged into one.

**Figure 1.**
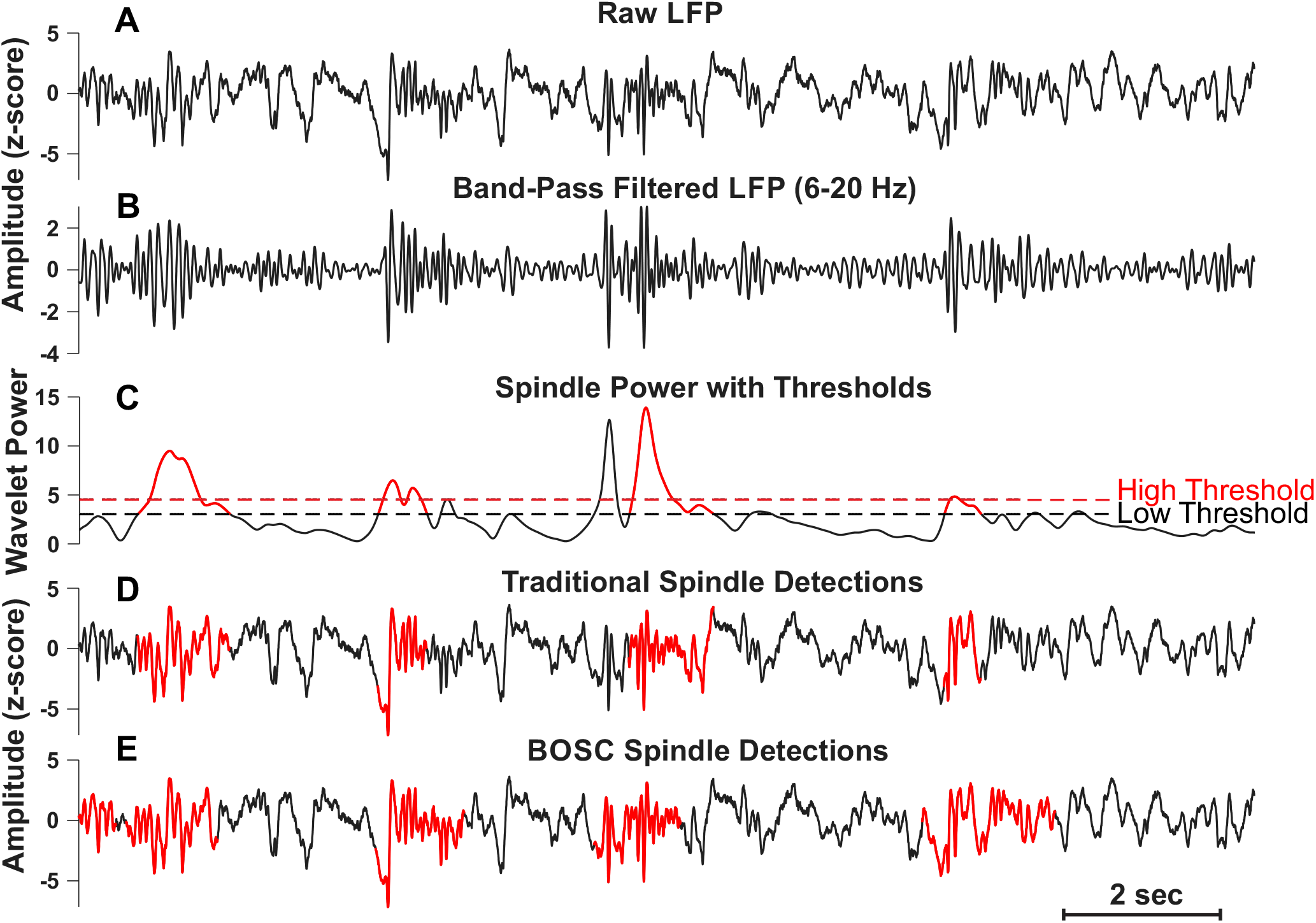
Overview of the traditional spindle detection workflow and comparison of detections with BOSC. (A) A raw local field potential (LFP/EEG) trace from rat frontal cortex during non-REM natural sleep. (B) A bandpass-filtered version of the same LFP segment within the spindle frequency range (6-20 Hz). (C) The Hilbert smoothed wavelet power within the spindle frequency range across time with power thresholds set by the traditional method (High=1.5 SD; Low=0.75 SD) and the detections it makes in red. (D) The raw LFP segment with the traditional spindle detections in red. (E) The raw LFP segment with the BOSC spindle detections in red.

### BOSC

We used a procedure based on that used by Witten et al. (2011). The BOSC method detects oscillatory episodes by identifying periods where the power at a specific frequency exceeds a power threshold — derived from the background frequency powers — for a minimum number of oscillatory cycles (“duration”; Fig. 2). The duration threshold set was scaled to each individual frequency. To analyze the signal, a spectral decomposition was performed using a Morlet wavelet transform (Grossmann & Morlet, 1985) with a width of 6 cycles and 21 logarithmically spaced frequencies ranging from 2 to 32 Hz. An estimate of the average background spectrum was made, assuming the signal has a colored noise model of the form *Af−^α^* (Schlesinger & West, 1988), which is characteristic of low-pass filtered autocorrelated signals. The resultant spectrum was fitted with a linear regression in log-log coordinates. A power threshold was set at the 99th percentile of the expected χ^2^(2) distribution of power values at each frequency of the background signal, using the mean power estimated from the linear regression. The χ^2^(2) distribution arises because Fourier-transforming a Gaussian-distributed LFP signal produces a complex number, whose squared magnitude (power) is a sum of two squared Gaussian components—one for the real and one for the imaginary part—resulting in a χ^2^ distribution with two degrees of freedom (Cox et al., 1993). The duration threshold was set to five full cycles of the oscillation (5/f). Oscillatory episodes were identified when both the power and duration thresholds were exceeded (Fig. 2 A). The proportion of time in which oscillations occurred at a given frequency, known as *P*_*episode*_, and the fit of the linear regression were used to evaluate and ensure the method’s performance on each given signal. Last, deviating from BOSC in its previously published implementation, any detections within 0.1 seconds of each other were merged into — one as is done in the traditional method.

**Figure 2.**
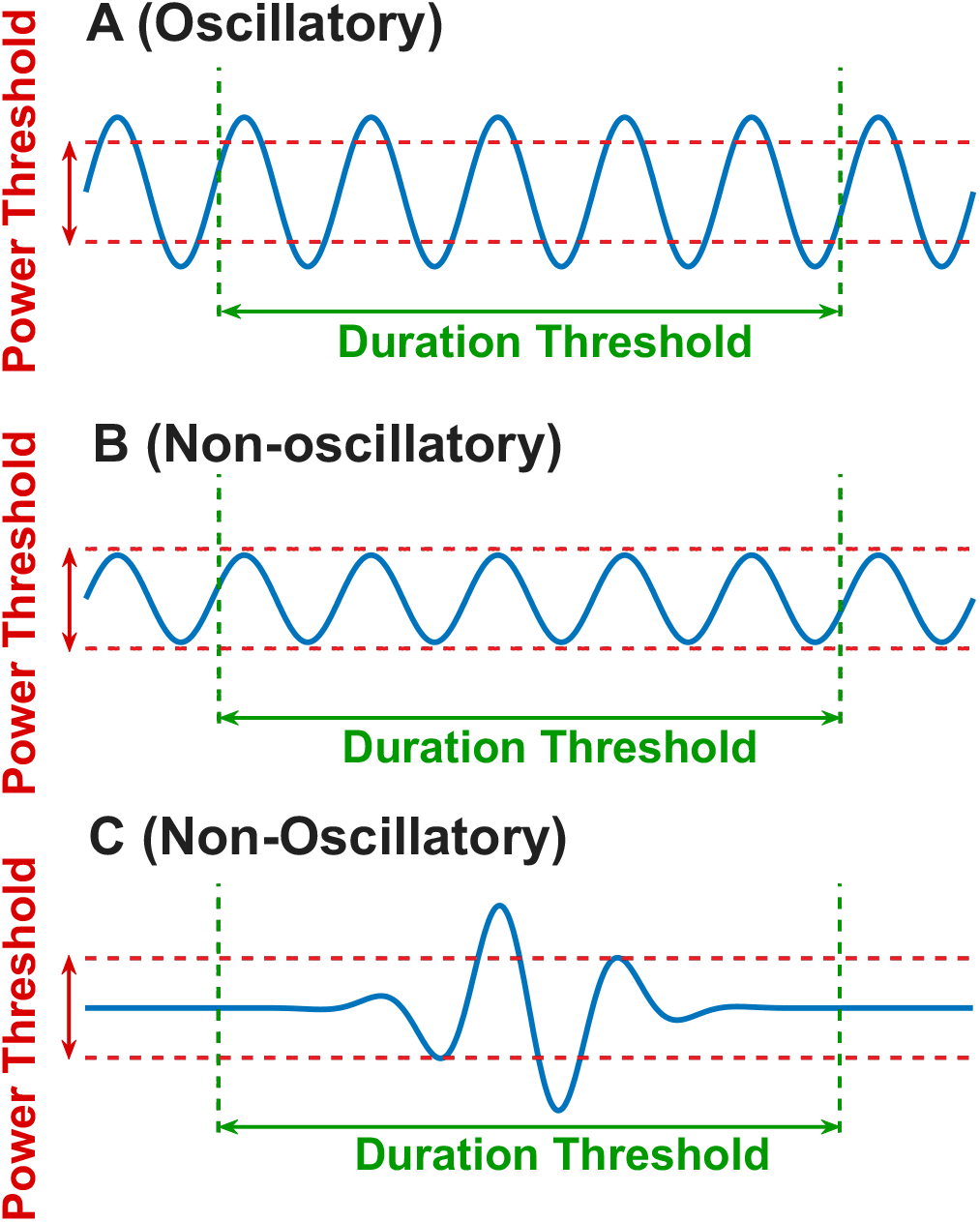
BOSC detection criteria (A) Example of an oscillatory event that exceeds both the power and duration thresholds required for BOSC to make a detection. (B) An oscillatory event that exceeds BOSC’s duration threshold but not the power threshold. (C) A non-oscillatory event that exceeds BOSC’s power threshold but not the duration threshold.

### Artificial LFP Signal Generation

The background signal used to add spindle-like activities was created from Gaussian noise restricted to amplitudes between ±1.00 (arbitrary units) that displayed a 1/f power spectrum. Noise of this nature was chosen because it mimics the frequency distribution of signals during natural sleep while maintaining an entirely random character. Because of its randomness, this allowed us to insert spindle-like activity into a situation with little to no expected periods of rhythmicity. This feature of the background signal was key in testing the abilities of our detection methods since it is relatively unlikely that spindle-like activity would arise in this signal. To create this synthetic 1/f noise, we constructed a frequency vector spanning 0.1–500 Hz and assigned each frequency an amplitude inversely proportional to its frequency. To better approximate natural LFP spectra, we applied a gradual suppression to the lowest frequencies (0.1-4 Hz) using a linear ramp to flatten the power slope below 4 Hz. Random variability was introduced by modulating the amplitude spectrum with bounded uniform noise (range: [-1, 1]),and phases were randomized across all frequencies. Last, an inverse fast Fourier transform was performed on the spectrum to obtain the time domain signal. The power spectral density curve of this signal was comparable to that recorded from the neocortex of a naturally sleeping rat (Fig. 3 A).

**Figure 3.**
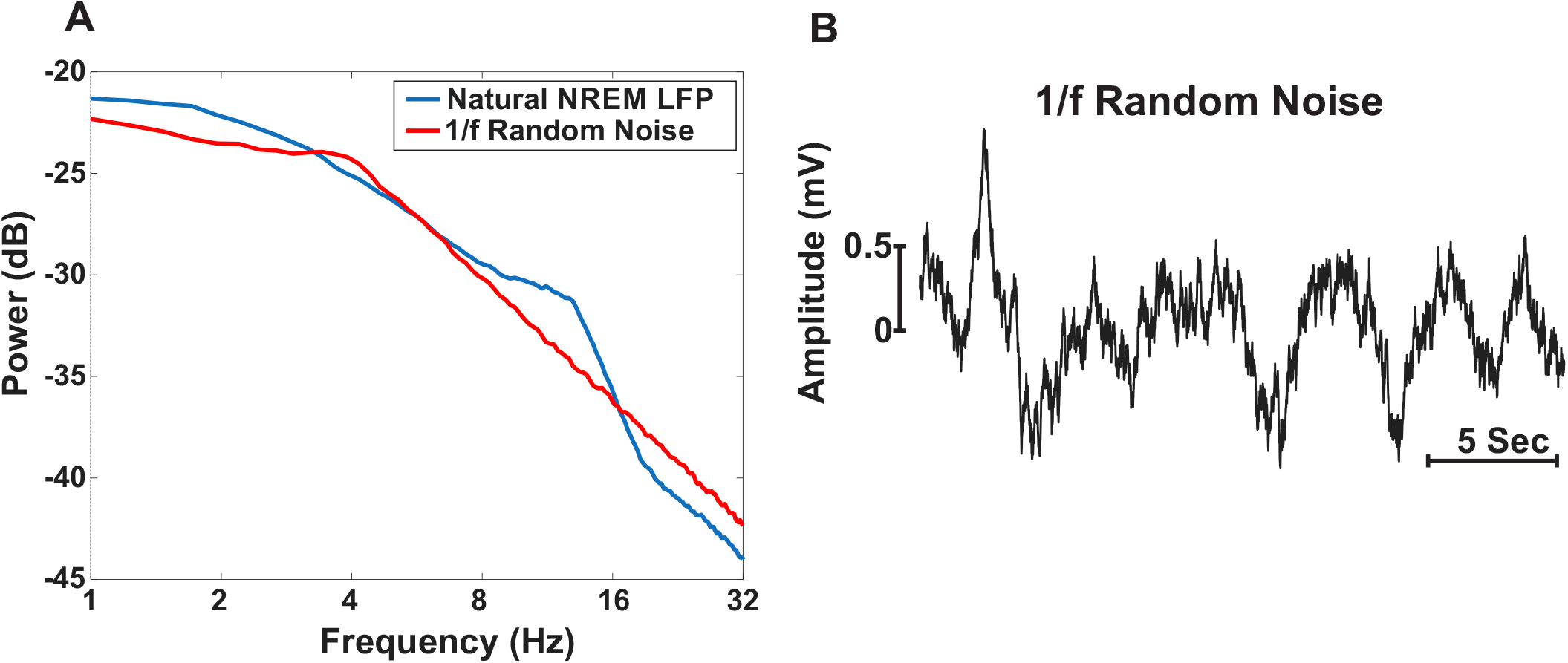
Comparison of synthetic 1/f background noise and natural sleep EEG (A) Power spectral density curves of the combined total NREM sleep from cortical recordings of a naturally sleeping rat in blue and random 1/f noise in red. Both curves follow a similar trend with the exception of the blue traces’ increase in power around the spindle frequency range (peak at ∽12 Hz). Such similarity justifies the use of random 1/f noise to model background NREM activity. (B) A representative segment of the 1/f random noise.

We then created synthetic and periodically occurring high voltage (HVS) and low voltage spindles (LVS) to add into our trace. We first synthesized a repeating sine wave with a consistent amplitude of ±1 (AU) that had wave-by-wave variations of its period that still fell within the frequency bandwidth of actual spindles (6-10 Hz for HVS and 10-20 Hz for LVS) (Johnson et al. 2010) (Fig. 4 A). This waveform was windowed using convolution procedures with a set of Gaussian functions applied throughout the time frame of the complex sine wave (Fig. 4 B-C). To mimic natural spindle amplitudes, the peak amplitude of each Gaussian window was sampled randomly from a distribution of actual spindle amplitudes observed and detected (using the traditional method) from rat recordings during NREM sleep (Fig. 5).^*^ When assigning amplitudes from the probability distribution, LVS were sampled using the distribution below the mean and HVS were sampled using the distribution above the mean. The series of synthetic spindles produced were then directly added to the random noise background signal at a specific period (every 50 seconds) with both HVS and LVS spindles interleaved (Fig 4. D-E).

**Figure 4.**
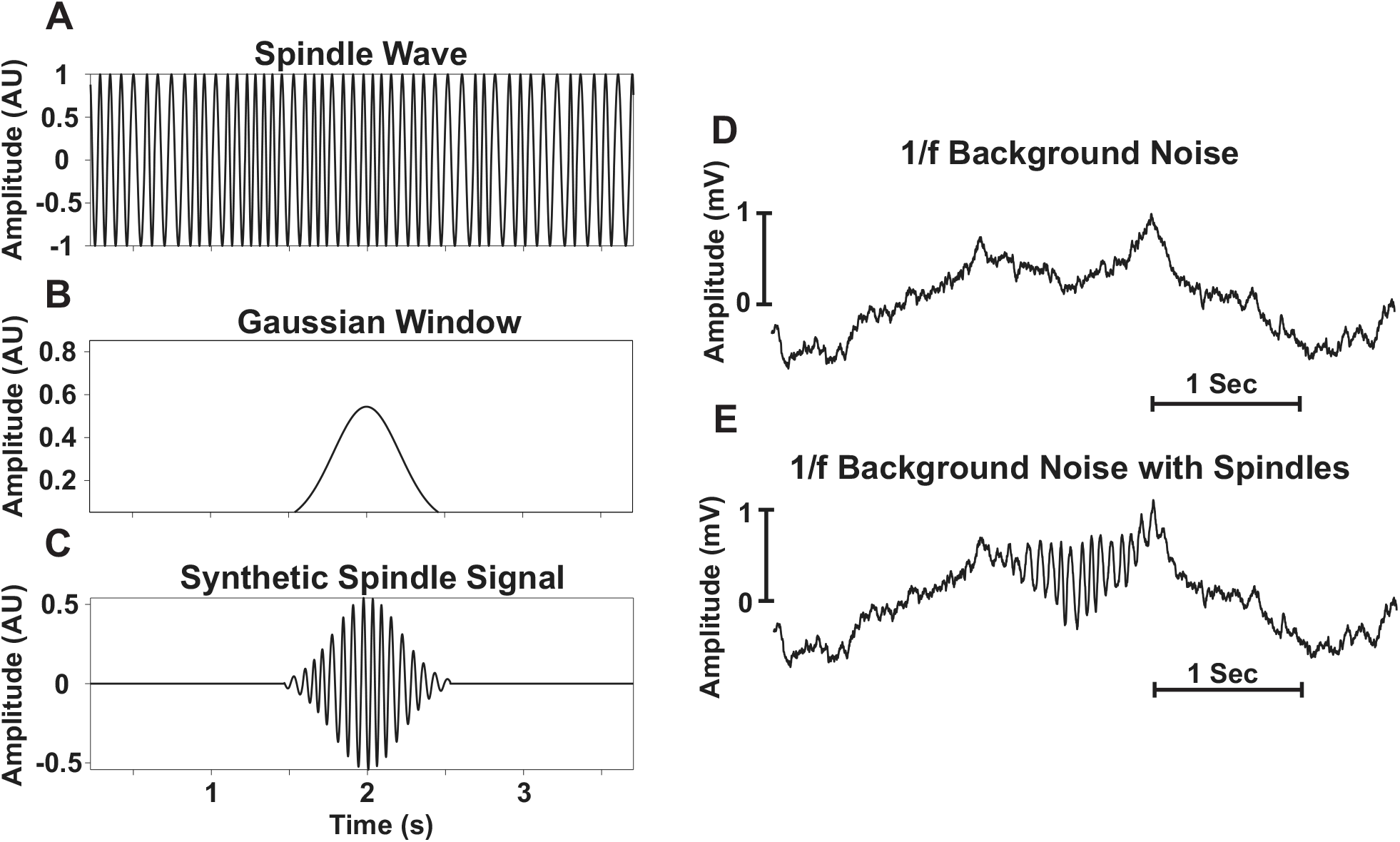
Construction of synthetic spindles and insertion into random 1/f background noise. (A) A segment of a generated signal composed of sine waves that show random variations in period corresponding to frequencies within the LVS frequency range (10-20) on a wave-by-wave basis. (B) An example of a Gaussian window function with an amplitude sampled from the probability density function in Figure 4 and with a random duration between 0.5 and 2 seconds. (C) The synthetic spindle signal produced upon convolutions of the spindle wave (A) and the Gaussian window (B). (D) A sample 1/f background segment with no added spindles. (E) The same background segment with the addition of the synthetic spindle signal. This final signal thus models the spectrum of natural NREM sleep with synthetic spindles added at regular intervals.

**Figure 5.**
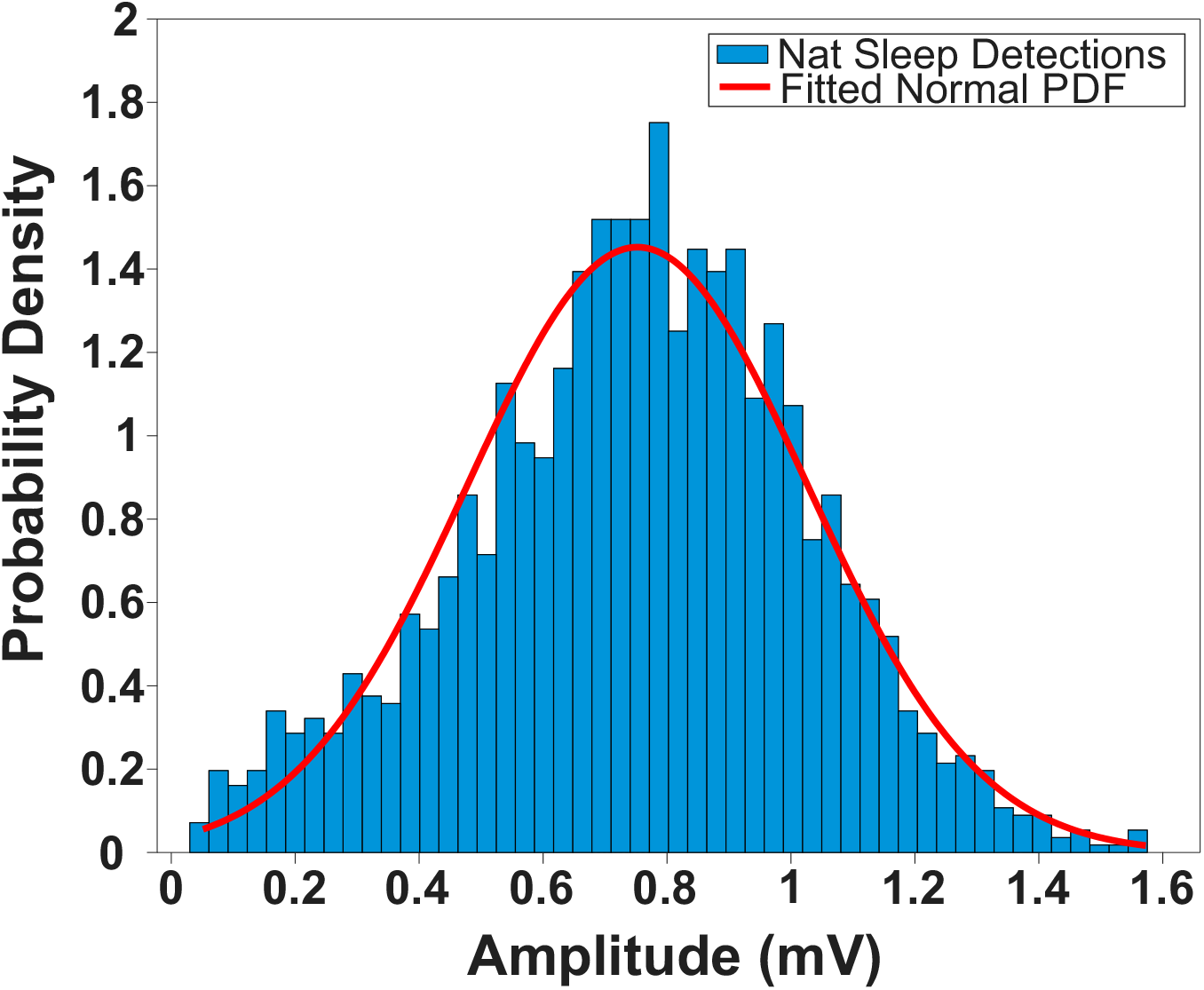
Generation of a probability density function from which to sample spindle amplitudes. A histogram showing the distribution of spindle amplitudes detected by the traditional method in a cortical LFP recording from a naturally sleeping rat. In red is a probability density function that shows a significant fit to the distribution (p = 0.1630, KS stat = 0.0262). This physiologically grounded distribution was used to sample amplitudes when generating synthetic spindles.

A similar method to the above was employed to add single pulses to our background noise. The only difference was that the duration of the Gaussian windows was restricted to 150 ms, and their amplitudes were set to produce power peaks between 0.5-2 AU. The period of these waves was also restricted to frequencies corresponding to a bandwidth of 6-8 Hz to ensure that no more than one cycle was possible (Fig. 6 A-C). This signal was then added to the same 1/f background as used for synthetic spindles above (Fig. 6 D-E)

**Figure 6.**
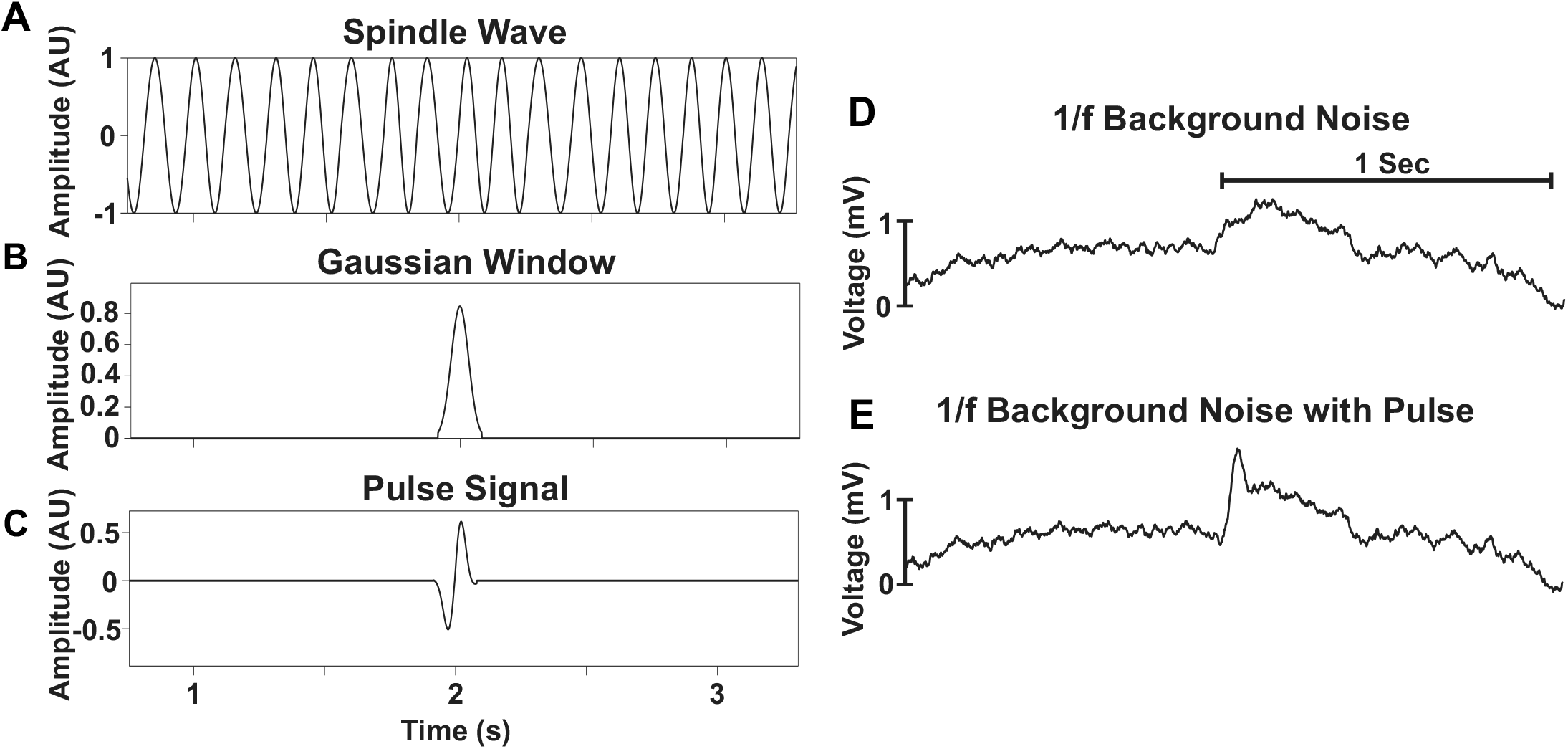
Construction of single wave pulses and insertion into random 1/f background noise. (A) A segment of a generated signal composed of sine waves that show random variations in period corresponding to frequencies within 6-8 Hz on a wave-by-wave basis. (B) An example of a Gaussian window function with an amplitude sampled from the probability density function in Figure 4 and with durations restricted to 150 ms to ensure that only one wave period was possible upon convolution. (C) The synthetic pulse signal produced upon convolution of the spindle wave (A) and the gaussian window (B). (D) A sample 1/f background segment with no added pulses. (E) The same background segment with the addition of the pulse signal. This signal models the background spectrum of natural NREM sleep with spindle frequency “spike” pulses added at regular intervals.

## Results

### Synthetic Spindle Detection

When testing the traditional method for spindle detections on our synthetic data, we found that it performed well. It correctly detected 91 out of 100 LVS, all 100 out of 100 HVS, and made 0 false detections. Testing BOSC in the same way using the same data, we found that it performed similarly. It correctly detected 94 out of 100 LVS, all 100 out of 100 HVS but with 2 false detections. Every detection made by the traditional method was also made by BOSC, but BOSC detected some unique events that included both true and false detections (Fig. 7). The F1 score, which represents the harmonic mean of precision (correct detections made / all detections made) and recall (correct detections made / all possible correct detections), was slightly higher for the BOSC method (F1 = 0.980) compared to the traditional method (F1 = 0.977), reflecting both a balance between accurate detection and avoidance of false positives in this dataset. Within each of these individual detections, the traditional method captured an average of 91.91 ± 1.56% of the true duration of each spindle while BOSC captured a lower average of 76.40 ± 1.36%.

**Figure 7.**
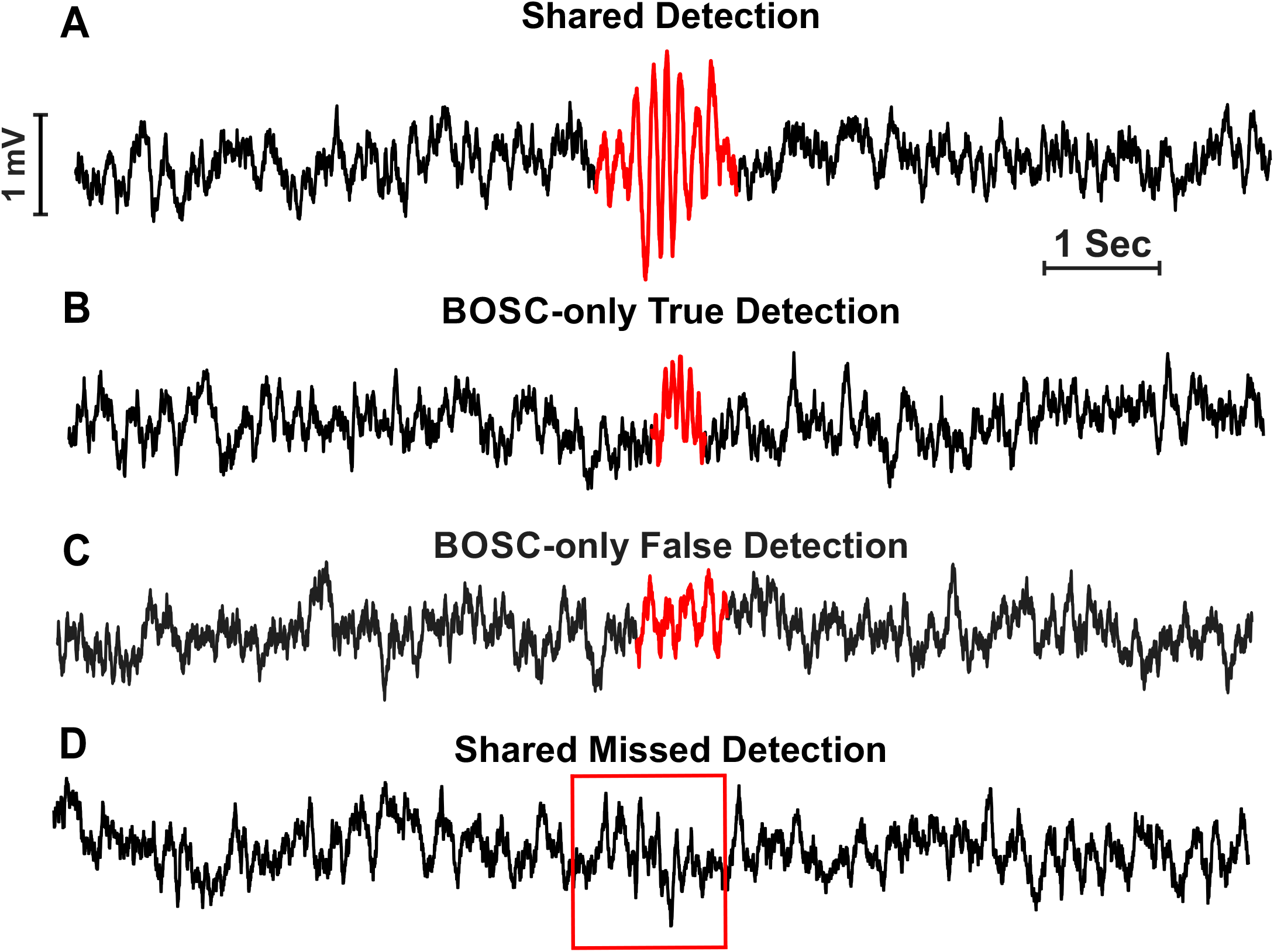
Performance of the traditional and BOSC methods on synthetic spindles. (A) Example of a synthetic spindle surrounded by random 1/f noise that was detected by both traditional and BOSC methods. (B) Example of a synthetic spindle that only BOSC detected. (C) Example of a 1/f background segment that was detected as a spindle by BOSC. (D) Example of a synthetic spindle that was not detected by either the traditional or BOSC method.

Since BOSC was making detections within the raw background trace, we tested how each method performed given only this 1/f noise. The traditional method made 657 detections including the 2 detections previously made by BOSC and BOSC made instead the same 2 detections as it did when the synthetic spindles were inserted.

### Synthetic One Wave Pulse Detection

Visual inspection of the false detections made by the traditional method when given the raw 1/f noise appeared to be related to high amplitude transient pulses. To quantify their susceptibility to detecting this noise, we tested how well each method handled the synthetic data with synthetic single wave pulses added. Any detections in this demonstration would be false positives. The traditional algorithm detected 94 out of 100 synthetic single wave pulses and made 135 other false detections. BOSC only detected 2 out of 100 single wave pulses and made the same 2 false background detections as with the synthetic spindles. The two single pulse detections made by BOSC were also made by the traditional method (Fig. 8 A). Interestingly, these detections did, in fact, resemble true spindles.

**Figure 8.**
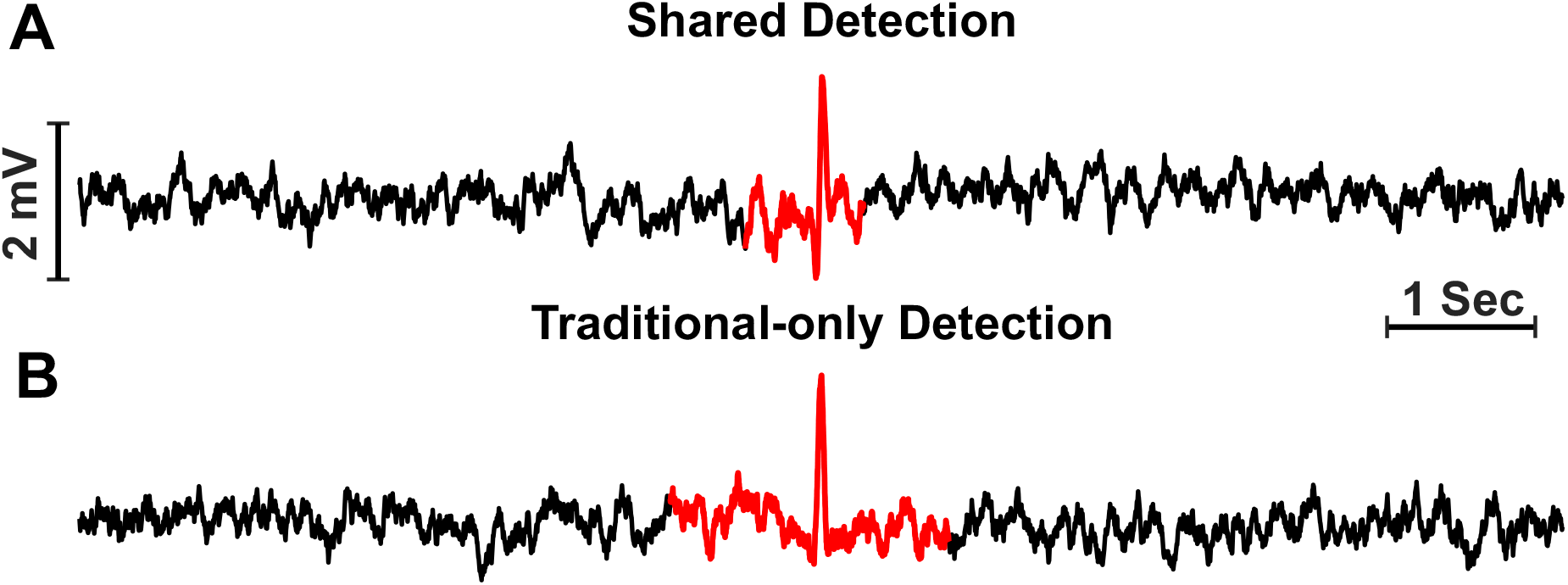
Performance of the traditional and BOSC methods on single wave pulses. (A) Example of a single wave pulse that was detected by both traditional and BOSC methods. (B) Example of a single wave pulse that was only detected by the traditional method. Every pulse that BOSC detected (2 out of the 100 inserted) was also detected by the traditional method.

### Natural Sleep LFP

To compare detections using natural sleep recordings, we analyzed cortical signals from 4 natural sleep experiments in rats using both the traditional and BOSC algorithms. The BOSC method detected more spindles by a factor of 1.59 (BOSC: 11979 detections, Traditional: 7479 detections). There were 5319 detections unique to the BOSC method, and 226 unique to the traditional method. Upon basic visual inspection of the detections that each method made (Fig. 9), it appeared that the detections each method had in common (A) and those unique to BOSC (B) displayed the typical increase in amplitude and waxing and waning shape expected of spindles while those unique to the traditional method (C) tended to be high amplitude pulses within the spindle frequency range, and as such, less convincingly spindles.

**Figure 9.**
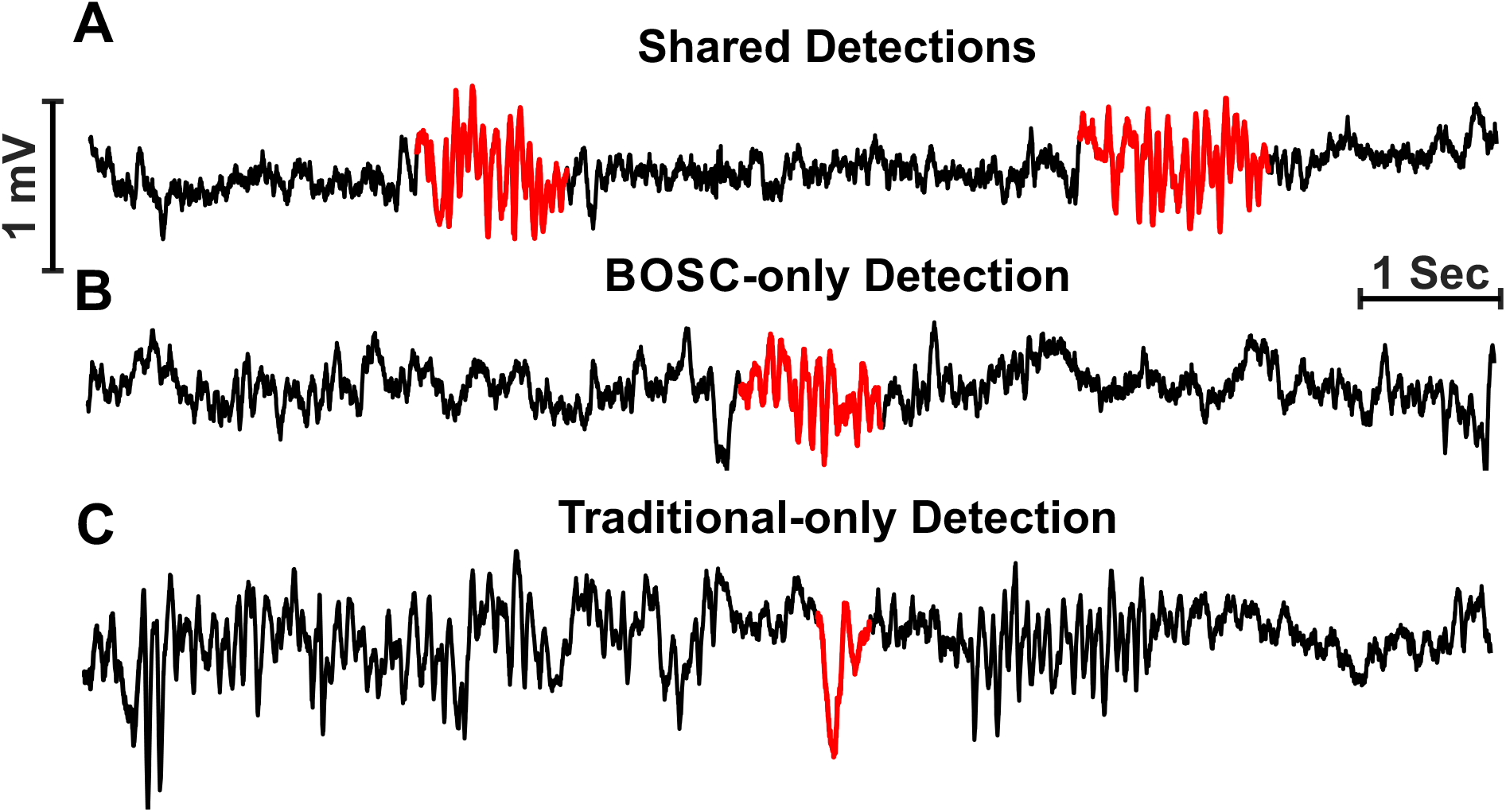
Performance of the traditional and BOSC methods on cortical LFP recordings from naturally sleeping rats. (A) Examples of overlapping spindle detections made by both traditional and BOSC methods. (B) Example of a spindle that only BOSC detected. Note the similarities to those in (A). (C) Example of a detection that only the traditional method made. Note the pulse-like nature of the event. These examples show how the BOSC tends to detect valid spindle activity that can be missed by the traditional method, while the traditional method can detect non-spindle activity.

We further confirmed the quality of detections for both methods by comparing the distributions of frequency, duration, and amplitude for each (Fig. 10). Both the traditional and BOSC methods showed similar distributions for each measure. For frequency (Column 1, Red), the distributions were highly overlapping and with statistically similar means (9.38-9.46 Hz, p > 0.05 for one-tailed paired t-test comparisons). However, BOSC had a slightly more pronounced bimodal frequency distribution. The distribution of spindle durations (Column 2, Green) showed modest differences, with BOSC showing a peak at a higher duration value than the traditional method and also showing a significantly higher mean value (BOSC: 1.72s, Traditional: 1.20s, one-tailed paired t-test, p < 0.001). Traditional-only durations also had a significantly lower mean value than the complete traditional duration distribution (Traditional-only: 0.52 s, Complete traditional:1.20 s, one-tailed paired t-test, p < 0.001). The amplitude distributions all resembled a positive-skewed normal distribution with the exception of the traditional-only case due to its lower number of events. Traditional spindles showed a significantly higher mean amplitude than that of BOSC (Traditional: 4.21 mV, BOSC: 3.72 mV, one-tailed paired t-test, p < 0.001), and the distribution of traditional-only amplitudes was also significantly higher than the BOSC-only amplitudes (Traditional: 3.78 mV, BOSC: 2.99 mV, one-tailed paired t-test, p < 0.001).

**Figure 10.**
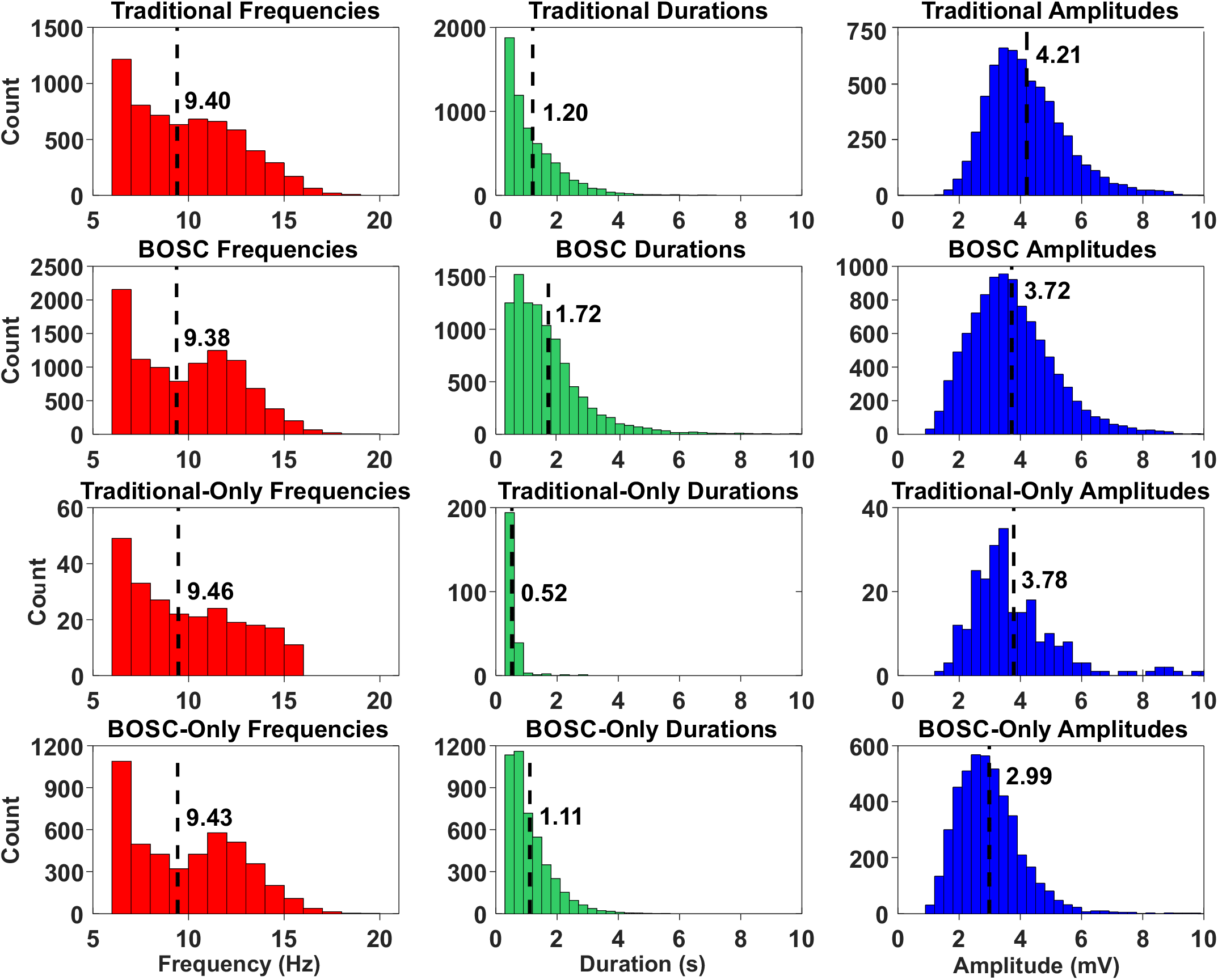
Spindle characteristics across detection methods. The first column (red) shows the distributions of all spindle frequencies detected from rat cortical LFP during natural sleep using the traditional (1st row) and BOSC (2nd row) methods as well as those detected only by the traditional (3rd row) and BOSC (4th row) methods. The second column (green) shows the distributions of all spindle durations detected under the same conditions. The third column (blue) shows the distributions of all spindle amplitudes detected under the same conditions.

Next, we analyzed the detections across state transitions using a peri-time histogram (binned) (Wake to NREM, REM to NREM, NREM to Wake, and NREM to REM) (Fig. 11). Both methods produced detections that were biased to the NREM state and the adjacent transition periods which is consistent with findings on spindle occurrence (Eschenko et al., 2006). Suspicious detections (i.e., those within Wake or REM that at latencies longer than 60s from NREM transitions) were coded and summed for each method. The traditional algorithm had a 1.08% (63/5834) suspicious detection rate and BOSC had a 1.26% (116/9808) suspicious detection rate. These values were statistically similar (z = -0.5848, p = 0.5587).

**Figure 11.**
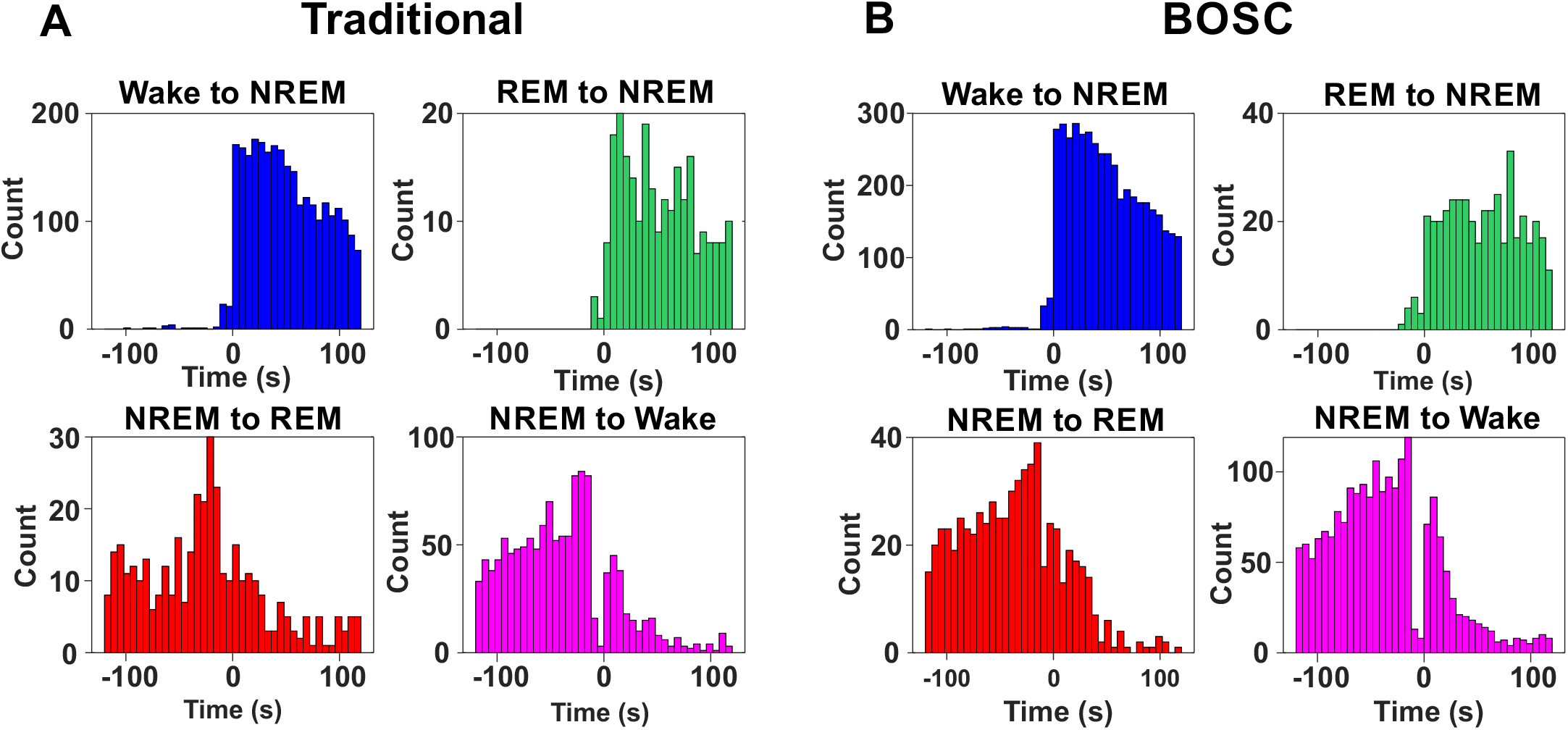
Temporal distribution of spindle detections around sleep state transitions. (A) Peri-event time histograms of spindle detections across four naturally sleeping rats using the traditional method around all four state transitions that involve NREM sleep (Wake to NREM, REM to NREM, NREM to REM, and NREM to Wake). (B) Peri-event time histograms of spindle detections for the same dataset and state transitions using the BOSC method. The general pattern of detections across these state transitions is conserved across the two methods.

## Discussion

Spindles are notoriously difficult to analyze due to their transient nature, complex circuitry and distribution across the cortex, as well as their potentially diverse functionality. Emerging research exploring spindles requires a robust detection method in order to ask the important functionally-related physiological questions; without one, the answers that follow may be in doubt. This research positions BOSC as a superior method of spindle detection compared to the traditional method due to its ability to avoid more non-oscillatory transients and detect more long duration and low amplitude spindles.

In terms of our comparisons of the two spindle detection methods here, our ground truth was based on the idea of generating synthetic spindles added into a random background signal with qualities comparable to that observed during NREM sleep in rats. In this way, we were able to truly test the abilities of both methods in a highly controlled fashion.

With the synthetic spindle traces, both methods performed almost equally well. For HVS, both the traditional and BOSC methods had a perfect detection rate. Although BOSC had slightly more correct detections of LVS, it also pulled out two “false” or “un-inserted” detections. In fairness to BOSC, the traditional method also made these same two detections when given the raw 1/f noise signal, along with 655 others, while BOSC still detected only the same two as before. This also shows that BOSC is less dependent on what is present in the signal of interest, whereas the traditional method performs differently at a given time in the recording depending on whether spindles are present or absent at other times in the recording. These “false” detections made by BOSC were visually reminiscent of low amplitude spindle events, and it is unclear how common events like these might be in random data such as ours (Fig. 7 C).

Furthermore, inspection of the 6 synthetic spindles that were not detected by either traditional or BOSC methods revealed that when added to the background signal, they appeared as short arhythmic low-amplitude events (Fig 7. D). Since these events met the criteria for spindles before insertion into the background signal it is clear that they were added in areas of the background that likely cancelled them out. Thus, the fact that they were omitted by both methods would actually be justified.

Another minor difference between the detection methods was the proportion of the actual spindle duration detected in these datasets, in which the traditional method performed better. This is both an advantage and a weakness of the traditional method based on the integration and smoothing provided by the Hilbert transform. Indeed, the tails of the synthetic signal have low amplitudes, and this is likely why they do not reach the power threshold required by BOSC. Additionally, this difference may not be relevant in the case of physiological spindles since BOSC still detects a broad spectrum of spindle durations that trend even longer in duration to those detected by the traditional method (Fig. 10).

Where the two methods completely diverged was in terms of the synthetic signals with single wave pulses added. Not only was the traditional method tricked by detecting practically all of these events (94%) it also made an even higher number of false detections, even beyond the original number of stimuli added. This is likely based on the relatively low average power within the spindle bandwidth and a subsequent lowering of the power threshold in the algorithm as a result. In contrast, the BOSC method was only fooled by 2 out of 100 pulses, which, when inspected, actually looked like true spindle activity based on the background activity at the time. BOSC’s superior performance on this task is likely attributed to its specified minimum oscillation requirement for detections.

Throughout the two synthetic dataset experiments, we tried to be as fair as possible to the traditional method by creating data that was not biased toward the strengths of BOSC. The first step was to use wavelet power for the traditional method in order to put it on more even footing with the wavelet analysis used by BOSC. They also included our use of the traditional method at the outset in order to create the distribution of amplitudes of spindles from rat neocortex, which then allowed us to create the synthetic spindles themselves. Also, we used randomized values for both frequency and duration of spindles derived from established ranges using the traditional method. We also used low-pass filtered 1/f random noise as our background because of its inherent stochasticity in order to create a realistic synthetic signal from which to test both methods equally.

When these two methods were compared in terms of detections of spindles in raw data, BOSC detected significantly more spindle activity (1.59 times more). Our characterization of the distributions of frequency, duration, and amplitudes for each of these detected spindles suggested first that the two methods detected spindles with properties that were highly similar and second, that the detections were likely valid spindles. The only potential exception was with the traditional method, since its detections that did not overlap with BOSC had properties that appeared almost exclusively at the shortest end of the duration distribution. Based on visual inspection, these events appeared as brief, non-oscillatory transients (spikes) or as noise misclassified as spindles based on high power in the spindle frequency range. In contrast, BOSC-only detections had similar distributions to its complete set, suggesting that they were true spindles that were simply undetected using the traditional method. BOSC detections also had a slightly more pronounced bimodal frequency distribution which likely distinguishes LVS from HVS (Johnson et al., 2010). Based on the differences between the various distributions across BOSC and the traditional method, we suggest that BOSC captures longer but lower-amplitude spindles that the traditional method overlooks, which is especially evident in the BOSC-only detections. These differences underscore BOSC’s advantage in detecting a more complete and correct dataset of spindle occurrences. As for why and how BOSC is able to capture this more complete and accurate representation of spindle activity, likely comes down to its minimum oscillation requirement which filters out non-oscillatory transients and it’s use of a fitted background power spectrum from which it derives the power thresholds. These thresholds can also be easily probed to suit the needs of the experimenter thus expanding BOSC’s use to other oscillatory contexts (Whitten et al., 2011).

That BOSC is able to detect longer duration and lower amplitude spindle activity has profound physiological relevance. Longer duration ripple-spindle interactions are of particular importance for memory consolidation, so increasing the temporal capabilities of a spindle detection method can give us a more complete picture of the key components involved in memory consolidation (Ngo et al., 2020). In addition, lower amplitude spindles are implicated in aging as well as various diseases and mental health conditions (Ferrarelli et al., 2007; Latreille et al., 2015; Martin et al., 2013). So, not only is the traditional method making less detections overall, but it is also excluding potentially physiologically relevant events.

In terms of further evaluating the accuracy of the two methods, both produced detections that were biased to the NREM state and the adjacent transition periods which suggests that the detections – like spindles – are restricted to NREM epochs. Worth noting is that during transitions from NREM to REM, both methods fail to directly cease detections upon the genesis of the REM state. This occurs because these transitions have a relatively long period that is characterized by significant bursts of spindle activity (Bandarabadi et al., 2020). In the case of the transition from NREM to Wake, there are also several detections in the earliest moments of the wake state which are – similarly – a result of a non-instantaneous transition period (Fernandez & Lüthi, 2020). Looking beyond the transition windows, both methods produced a similar number of suspicious detections during REM and Wake states which we hesitate to call false detections due to the often ambiguous and unstable nature of states when recording from a unique and precise cortical location during behaviour (Mohajerani et al., 2013; Niethard et al., 2018). Combined with the observation that both methods generally follow expected patterns of activity across state transitions, this suggests that, for the most part, traditional detections are not entirely invalid. However, based on the number of detections, and their individual qualities, it would seem like BOSC is much more sensitive and reliable.

## Conclusion

In this study, we compared the performance of the traditional power-threshold spindle detection method with the Better OSCillation detection (BOSC) algorithm using both synthetic and biological LFP data. Our findings demonstrate that BOSC offers a more sensitive and reliable approach to spindle detection. In synthetic datasets, BOSC and traditional methods reliably detected nearly all synthetic spindle events while avoiding false positives; however, the traditional method was highly unreliable (in contrast to BOSC) when presented with synthetic single spindle frequency pulses. BOSC’s advantages stem from its use of frequency-specific background estimation and a minimum cycle threshold, enabling it to distinguish genuine oscillatory activity from non-oscillatory transients. Furthermore, in actual neocortical recordings, BOSC identified many more events than the traditional method, and the latter detected unique events that were unlikely to be actual spindles. Our results underscore the utility of BOSC as a robust and principled method for spindle detection in both experimental and clinical LFP/EEG research. Its improved sensitivity to subtle but meaningful oscillatory activity may provide a more comprehensive window for the understanding of the role of sleep spindles in memory consolidation, cognitive development, and neuropsychiatric conditions. Future work can extend this approach to other transient oscillatory phenomena (eg. ripples) or apply it to the analysis of spindles under different experimental conditions.

## Acknowledgements

This work was supported by the Natural Science and Engineering Council of Canada (NSERC) discovery grant 2021-02926 to C.T.D. and 2021-02551 to S.P. An early draft of this manuscript was reviewed by Dr. Jeremy Caplan who provided helpful comments. The authors declare no competing financial interests.

## Data Repository

https://data.mendeley.com/datasets/hzw6zpzp9j/1

## Footnotes

* To determine what amplitudes to assign the synthetic spindles we used the traditional method to detect spindles from one of our natural sleep experiments and measured each detection′s amplitude by measuring the maximum peak to trough distance and dividing by 2 (Fig. 5). All measures less than 0.05 mV (less than 1% of the dataset) were removed. The advantage of this approach is that most of the background influence on amplitude is removed, and we are left with a reasonable estimation of actual spindle peak amplitudes. The fit of the probability distribution to the actual distribution of amplitudes was verified using a Kolmogorov-Smirnov test which suggested that the two distributions were not significantly different (p = 0.1630, D = 0.0262). We employed the traditional method here in order since it is already an accepted method of detection and we wanted to be as fair to its utility as possible.

## Notes

https://data.mendeley.com/datasets/hzw6zpzp9j/1

### Competing Interest Statement

The authors have declared no competing interest.

